# Unraveling and quantifying “*Candidatus* Saccharibacteria”: *in silico* and experimental evaluation of V3-V4 16S rRNA metagenomics and qPCR protocols

**DOI:** 10.1101/2024.03.06.583507

**Authors:** Stella Papaleo, Riccardo Nodari, Lodovico Sterzi, Enza D’Auria, Camilla Cattaneo, Giorgia Bettoni, Clara Bonaiti, Ella Pagliarini, Gianvincenzo Zuccotti, Simona Panelli, Francesco Comandatore

## Abstract

**Background:** Candidate Phyla Radiation (CPR) is a large monophyletic group thought to cover about 25% of bacterial diversity. Due to peculiar characteristics and unusual 16S rRNA gene structure, they are often under-represented or lost in 16S rRNA-based microbiota surveys. Among CPR, “*Candidatus* Saccharibacteria” is a phylum experimentally found to modulate the immune response and enriched in the oral microbiota of subjects suffering from several immune-mediated disorders, e.g. food allergies, as reported by us in a previous work. Due to the growing evidence of “*Ca*. Saccharibacteria”’s role in clinical settings and in order to unravel its role in host physiology and pathology, it is crucial to have a reliable method to detect and quantify this lineage.

**Methods and Results:** Four qPCR protocols for quantifying “*Ca.* Saccharibacteria” (one targeting the 23S rRNA gene and three the 16S) were selected from the literature among the few available. Efficiency and coverage of primer pairs used in these protocols were preliminary evaluated via *in silico* analyses on the “*Ca.* Saccharibacteria” known taxonomic variability, and then tested *in vitro* on the salivary DNA previously investigated by 16S metagenomics in the food allergy study. *In silico* analyses evidenced that the 23S qPCR protocol covered more “*Ca*. Saccharibacteria” variability compared to the 16S-based ones, and that the 16S metagenomics primers were the most comprehensive. qPCR experiments confirmed that 16S-based protocols strongly underestimated “*Ca*. Saccharibacteria” while the 23S protocol was the only one to yield results comparable to 16S metagenomics both in terms of correlation and absolute quantification. However, only 16S metagenomics evidenced an expansion of “*Ca*. Saccharibacteria” in allergic subjects compared to controls, while none of the four qPCR protocols detected it.

**Conclusion:** These results underline the current limits in experimentally approaching “*Ca*. Saccharibacteria”. To obtain a more realistic picture of their abundance within bacterial communities, and to enable more efficient taxonomic resolution, it is essential to find novel experimental strategies. This is a necessary premise for more targeted and systematic functional studies to clarify the role of “*Ca*. Saccharibacteria” and, generally, CPR bacteria, in maintaining the health of the host.

## 1. Introduction

In the last decades, culture-independent molecular methods allowed the discovery of a large new group of bacteria from environments and human bodies, now referred to as Candidate Phyla Radiation (CPR) (Torrella and Morita 1981; Brown et al. 2015; Hug et al. 2016; Castelle and Banfield 2018). Currently, this monophyletic bacterial lineage includes more than 70 phyla (Danczak et al. 2017; Naud et al. 2022) and is still called “candidate” due to the lack of cultivated representatives, except for a few exceptions (Murugkar et al. 2020; Ibrahim et al. 2021a). CPR population structure is currently poorly understood and the size of the CPR group is still debated. Recently, it has been estimated that it encompasses about 25% of the bacterial diversity (Nie et al. 2022).

CPR are small-sized cocci (0.2-0.3 μm) with reduced genome size (usually < 1 Mb) (Luef et al. 2015) lacking important pathways, as those for aminoacids and nucleotide biosynthesis (Brown et al. 2015). Shotgun metagenomics highlighted that they have an unusual ribosome composition, missing some ubiquitous bacterial genes, such as uL1, bL9, and/or uL30. Furthermore, they have a peculiar 16S rRNA gene sequence with introns and indels (Tsurumaki et al. 2022). The few successful cultivation attempts led to the discovery of unique lifestyles, with CPR colonizing the surface of other bacteria within the community, and living as epibionts with mutualistic/parasitic lifestyles (Gong et al. 2014; He et al. 2015).

CPR phyla as “*Candidatus* Saccharibacteria” (formerly known as TM7), “*Candidatus* Absconditabacteria” (SR1) and “*Candidatus* Gracilibacteria” (GNO1) are now considered as part of the microbiota of human healthy oral tract, stomach and skin. Furthermore, either observational and experimental studies converged in suggesting their medical importance (Bor et al. 2019); (Naud et al. 2022).

Among these lineages, “*Ca.* Saccharibacteria’’ is the most studied. It has been reported to represent at least 3% of the human core oral microbiota and to be enriched in dysbiotic microbiomes during infection and inflammatory states of the oral mucosa (e.g., periodontitis and gingivitis, (Bor et al. 2019), and beyond (i.e., in Inflammatory Bowel Disease patients, Naud et al. 2022). These bacteria live as obligate epibionts (either mutualistic or parasitic), colonizing the surface of *Actinobacteria*, a phylum of bacteria usually present in human oral microbiota. The *Actinobacteria* host can belong to species with the potential to cause proinflammatory effects to the human counterpart. The epibiont can in turn modulate these inflammatory effects and have immunomodulatory activities itself on the human host (He et al. 2015; Chipashvili et al. 2021). These effects have been studied on *Nanosynbacter lyticus* (previously, TM7x), the first lineage within “*Ca.* Saccharibacteria’’, and the first CPR, to be isolated in coculture with its host, *Actinomyces odontolyticus* (now *Schaalia odontolytica)* strain XH001 (He et al. 2015). *S. odontolytica* has a strong pro-inflammatory effect by inducing Tumor Necrosis Factor Alpha (TNF-α) gene expression in macrophages. *N. lyticus* is able to suppress TNF-α expression and to prevent the detection of its host by human macrophages (He et al. 2015). This anti-inflammatory effect of *N. lyticus*, as well as of other “*Ca.* Saccharibacteria” species isolated in coculture in the meanwhile, have been confirmed by subsequent functional studies (Chipashvili et al. 2021).

Due to the growing awareness of its clinical relevance, it is important to have reliable methods to detect and quantify “*Ca.* Saccharibacteria” in human microbiota in various physiological and pathological conditions. This is a necessary premise for more focused taxonomic and functional studies, to clarify their population structure and role in maintaining the host’s health status. Unfortunately, given the peculiar characteristics of CPR bacteria, current molecular methods work poorly on them, or give biased pictures, especially regarding the estimate of relative abundances. As regards the amplicon sequencing, the most frequently used “universal” primers on the 16S gene display a low efficiency in amplifying CPR sequences (Brown et al. 2015; Eloe-Fadrosh et al. 2016). On the other hand, in the last years several qPCR protocols targeting 16S or 23S rRNA genes have been designed for the quantification of “*Ca.* Saccharibacteria” in various environments (Takenaka et al. 2018a; Ibrahim et al. 2021b).

In this work, we evaluated four published qPCR protocols for “*Ca.* Saccharibacteria”, three designed on 16S and one on 23S rRNA gene. An *in silico* analysis was firstly performed on sequences representative of the whole known taxonomic variability within “*Ca.* Saccharibacteria”. qPCR experiments were then performed using the same salivary DNA samples from children suffering from food allergy and matched controls, previously characterized by us using the V3-V4 16S metagenomics (D’Auria et al. 2023). In that previous work, the oral microbiota of allergic children was found to be enriched in “*Ca.* Saccharibacteria” and unclassified bacteria. Here, we reevaluated the presence and relative abundance of “*Ca.* Saccharibacteria” in these samples, also in the light of the *in silico* analyses, to get more insights into the drawbacks and distortions associated with the currently available protocols for detecting, quantifying and classifying this emerging bacterial lineage.

## 2. Materials and Methods

### 2.1. Selection of primer pairs

The current efficiency in the detection and quantification of “*Ca.* Saccharibacteria’’ was assessed *in silico* and through qPCR experiments using six primer pairs retrieved from the literature (see Table S1): SacchariF-SacchariR (Ibrahim et al. 2021a) (here called 23S), TM7314F-TM7-910R (Hugenholtz et al. 2001; Brinig et al. 2003) (16S_p1), Sac1031F-Sac1218R (Yang et al. 2015) (16S_p2), TM7_16S_590F-TM7_16S_965R (Ferrari et al. 2014) (16S_p3), 926F-1062R (Bacchetti De Gregoris et al. 2011) (16S_panbacteria), and pro314F-pro805R (Takahashi et al. 2014) (16s_meta).

The latter are V3-V4 primers commonly used in 16S rRNA metagenomic studies. We used this pair in our previous work on the salivary microbiota of allergic children (D’Auria et al., 2023). The other pairs have been designed for qPCR and were included in the present study for the reasons detailed below.

The 23S protocol was chosen because primers are based on a very recent genomic analysis ((Takenaka et al. 2018a; Ibrahim et al. 2021b) and because it targets a gene other than the 16S rRNA, known to have a limited capacity to detect CPR.

Two out of the three 16S rRNA primer pairs (protocols 16S_p1 and 16S_p2) were chosen based on Takenaka et al. (2018) (Takenaka et al. 2018a; Ibrahim et al. 2021b) that evaluated different primers for “*Ca.* Saccharibacteria’’ quantification. These authors concluded that TM7314F/TM7-910R (16S_p1) gave the most reliable real time quantification, and for this reason we included them in our collection. The other pair, Sac1031-F/Sac1218R (16S_p2) in their hands appeared to underestimate their environmental samples, but because it was originally designed to analyze “*Ca.* Saccharibacteria’’ in mammalian feces (Yang et al. 2015) we decided to test it on our dataset. The third 16S pair (TM7_16S_590F/TM7_16S_965R, protocol 16S_p3), described in (Ferrari et al. 2014), has already been recognized for its high coverage and specificity for “*Ca.* Saccharibacteria” (Takenaka et al. 2018b).

The 926F-1062R pair (16S_panbacteria) (Bacchetti De Gregoris et al. 2011) is a pair of universal 16S primers commonly used for qPCR. It was used in combination with the “*Ca.* Saccharibacteria” pairs to evaluate by qPCR their abundance in reference to the total bacterial quantification.

### 2.2. In-silico PCR experiments

The efficiency of the primer pairs listed above was first tested *in silico* PCR on two large datasets: the “*Candidatus* Saccharibacteria” sequences contained in the SILVA database and a collection of high quality “*Candidatus* Saccharibacteria” genomes. Regarding the SILVA database, the reference datasets LSU Ref NR99 v.138.1 (Large Subunit, i.e. 23S rRNA gene) and SSU Ref NR99 v.138.1 (Small Subunit i.e. 16S rRNA gene) were retrieved and the sequences annotated as “Saccharimonadia” (the only SILVA annotation relative to Saccharibacteria) were extracted. Unfortunately, only two “Saccharimonadia” sequences were present in the LSU Ref NR99 v.138.1 (i.e. 23S rRNA gene), and thus the *in silico* PCR analyses could be carried out only on the SSU Ref NR99 v.138.1 (16S rRNA gene) dataset, from which 2,978 “Saccharimonadia” sequences were extracted. The *in silico* PCR analyses were performed using the ThermonucleotideBLAST tool (Gans and Wolinsky 2008) setting the following parameters: --primer-clamp 5 --max-mismatch 6 --best-match -m 1.

The 2,978 extracted sequences were then aligned using the MAFFT tool (Gans and Wolinsky 2008) and phylogenetic analysis carried out using FastTree (Price, Dehal, and Arkin 2010). The results of the *in silico* PCR were mapped on the obtained phylogenetic tree using iTOL web tool (Price, Dehal, and Arkin 2010; Letunic and Bork 2021).

The same analysis was repeated on the 16S rRNA and 23S rRNA gene sequences of a second large dataset, a manually curated collection of “*Candidatus* Saccharibacteria” genomes, as follows. All the “*Ca.* Saccharibacteria” genome assemblies present into the BV-BRC database (Price, Dehal, and Arkin 2010; Letunic and Bork 2021; Olson et al. 2023) as of June 27, 2023 were retrieved and subjected to 16S rRNA and 23S rRNA gene calling using Barrnap (github.com/tseemann/barrnap). The 16S rRNA sequences sized between 1,300 and 1,500 nt, and the 23S rRNA sequences sized between 3,000 and 3,500 nt, were considered complete. The genome assemblies harboring at least one complete 16S rRNA and one complete 23S rRNA gene were selected. For each genome, all the 16S rRNA gene sequences called by Barrnap were analyzed by *in silico* PCR as described above, using the five primer pairs targeting 16S (16S_p1, 16S_p2, 16S_p3, 16S_panbacteria and 16S_meta primers); the same was done for the 23S rRNA gene and the corresponding primer pair. The longest 16S rRNA sequence of each selected genome was extracted and subjected to phylogenetic analysis using FastTree, after alignment using MAFFT. The results of the five *in silico* PCR experiments (five on 16S rRNA gene target and one on the 23S rRNA gene) were mapped on the obtained phylogenetic tree using iTOL (Letunic and Bork 2021).

As described below, *in vitro* experiments were carried out on DNA previously extracted and subjected to 16S metagenomics by D’Auria et al (D’Auria et al. 2023). The V3-V4 16S rRNA sequences annotated as Saccharibacteria by D’Auria and colleagues (2023) were retrieved and Blastn searched against both the two 16S rRNA datasets (from SILVA and genome assemblies) already used for phylogenetic analyses. For each sequence, the most similar sequence was highlighted on the phylogenetic trees using iTOL (Letunic and Bork 2021).

### 2.3. DNAs and Primers

The four qPCR protocols for quantification of “*Ca*. Saccharibacteria” were tested *in vitro* on 61 DNA samples already subjected to 16S metagenomics by D’Auria et al (D’Auria et al. 2023). In that study, DNA was extracted from saliva of patients suffering from food allergies and matched controls, and subjected to 16S metagenomics. The same DNA preparations were used in this study: samples were not re-extracted in order to avoid any kind of variation that would have distorted the comparison between the qPCR results and the metagenomic analysis. The quantifications obtained with the pairs targeting “*Ca.* Saccharibacteria” (16S_1, 16S_2, 16S_3 and 23S) were normalized on the total bacterial DNA quantification of the sample, performed with 926F-1062R (here called 16S_panbacteria), a pair of universal 16S primers commonly used for qPCR (Bacchetti De Gregoris et al. 2011).

### 2.4. PCR protocols

For each primer pair, a standard end-point PCR protocol was first run to verify specificity and provide amplicons for the standard curve for subsequent qPCR experiments. PCR reactions were performed on those salivary DNA samples that, following amplicon metagenomics, displayed the highest relative abundances of “*Ca.* Saccharibacteria”. Amplifications were set up in a total volume of 20 μL containing: 10 μL GoTaq® Green Master Mix (Promega Corporation, Madison, Wisconsin, USA), 1 μL of each 10 μM primer, 6μL Promega PCR amplification-grade water (Promega) and 2 μL of the sample DNA (corresponding to about 20 ng). Cycling programs were performed on a Biorad T100 thermal cycler. Thermal profiles are listed in Table S2. PCR products were analyzed through electrophoresis on 1% agarose gels. Amplicons were gel-purified using Wizard® SV Gel and PCR Clean-Up System (Promega) and quantified with a Qubit 4 Fluorometer (Thermofisher scientific, Waltham, Massachusetts). DNA was finally diluted in Milli-Q water. Ten-fold serial dilutions were prepared for each amplicon that contained known numbers of fragment copies ranging from 10^7 to 10 copies/μL to create the standard curves.

### 2.5. qPCR protocols

Each 15 μL reaction contained 7,5 μL of 2x SsoAdvanced Universal SYBR® Green Supermix (BioRad, Hercules, California), 0,4 μL of each 10 μM primer, 4,7 μL of PCR amplification-grade water (Promega Corporation, Wisconsin, USA) and 2 μL of sample DNA (about 20 ng). Each sample was qPCR-amplified in three technical replicates. The qPCR assays were performed on a BioRad CFX Connect real-time PCR System (BioRad, Hercules). Thermal profiles are listed in Table S3.

The specificity of each primer pair was assessed through the melting profile generated at the end of each qPCR experiment, with a range of temperature between 60° and 95°C.

### 2.6. Statistical analyses

The detecting capability of the four primer sets tested in this study (16S_p1, 16S_p2, 16S_p3 and 23S) was compared on the basis of the “*Ca.* Saccharibacteria” quantification provided by each of them, as follows. For each primer set, the “*Ca.* Saccharibacteria” representation in the total bacterial community was calculated, in percentage, as the ratio between their absolute quantification and the pan-bacterial absolute quantification obtained using the 16S_panbacterial primers (see Table S1-S3 for details). Results obtained in this way for each of the four qPCR primer sets, and those from the 16S metagenomics (D’Auria et al. 2023), were then compared with Mann-Whitney U test and linear regression (significant p value threshold 0.05), using R. For each of these five methods of quantification, the “*Candidatus* Saccharibacteria” percentages obtained for allergic vs control subjects were compared using Mann-Whitney U test, using R.

### 2.7. Sequencing and analysis of 23S rRNA gene amplicon

Twelve representative samples selected from the 61 tested first by 16S metagenomics (D’Auria et al., 2023) and then by qPCR were chosen for 23S amplicon sequencing, to verify the specificity of the primers and define the portion of the taxonomic variability of “*Ca*. Saccharibacteria” covered by these primers. Eight samples were chosen because they displayed the highest differences between the quantifications provided by the 23S qPCR and those obtained from the 16S metagenomics, while other four samples were sequenced as controls. Sequences were performed on an Illumina Novaseq 6000 platform by MrDNA, Shallowater, Texas. Reads quality was assessed using the FastQC tool (http://www.bioinformatics.babraham.ac.uk/projects/fastqc). Then, the 23S rRNA gene amplicon reads were taxonomically assigned using the Mothur tool (Schloss et al. 2009) and SILVA138.1 LSURef NR99 as reference database (Quast et al. 2013). Briefly, reads were aligned against the reference Silva database and those containing chimeric information were removed. The remaining reads were grouped into Operative Taxonomic Units (OTUs) using the 0.05 distance threshold (without *a priori* information, the threshold has been determined on the basis of the nucleotide distance distribution). Then, a phylogenetic-based taxonomic annotation of OTUs was performed on the representative reads of the different OTUs. The reads were BlastN-searched against the NCBI nt database and, for the 20 best hits, sequences and taxonomic metadata were retrieved. The obtained NCBI sequences and the representative OTU sequences were aligned and subjected to Maximum Likelihood (ML) phylogenetic analysis using RAxML8 (Stamatakis 2014), with 100 pseudo bootstraps, using the model K80+G, as determined by best model selection analysis using ModelTest-NG (Darriba et al. 2019).

## 3. Results and discussion

### 3.1. In silico PCR experiments

*In silico* PCR analyses were performed on sequences representative of the whole known taxonomic variability within “*Ca*. Saccharibacteria”, retrieved from two large datasets. These sequences are the 2,978 16S rRNA annotated as Saccharimonadia retrieved in the SILVA database (Quast et al. 2013), and the 16S/23S rRNA sequences from a manually curated 114 “*Ca*. Saccharibacteria’’ genomes dataset (Table S4).

Figure 1 shows the 16S rRNA-based phylogenetic trees obtained for the two datasets (hereafter referred to as “SILVA” and “genomes”), annotated with the results of the *in silico* PCR analyses for all the six sets of primers considered in this study (see Methods and Table S1 for details). The colored rings in Figure 1 indicate the taxonomic variability within “*Ca*. Saccharibacteria’’ successfully amplified by each pair. Results for SILVA (Figure 1a) evidenced that none of the protocols completely covered the taxonomic variability. The highest coverage was obtained for 16S_meta, i.e., the primers for 16S metagenomics, that *in silico* amplified 97.5% (2,903) of the 2,978 “Saccharimonadia’’ 16S rRNA sequences in SILVA. Similarly, 96.5% of the sequences (2,875) was amplified by 16S_panbacteria primers, followed by 83.3% for 16S_p3 (2,482), 64% (1,908) for 16S_p1, and only 5.6% (168) for 16S_p2. As explained above (see Methods, paragraph 2.2) this analysis could not include the 23S primers because of the poor representation of 23S rRNA sequences belonging to “*Ca*. Saccharibacteria” in the SILVA database. Overall, the results for SILVA showed that, in this quite large dataset, all *in silico* amplifications missed a variable portion of the currently known taxonomic variability within “*Ca.* Saccharibacteria” (probably far from exhaustive), with the best “performance” highlighted for the 16S_meta pair, which missed the 2.5% of Saccharimonadia 16S rRNA sequences. This suggests that, even though 16S_meta primers have a very high coverage for the “*Ca.* Saccharibacteria” phylum (which is the one, within the CPR group, for which most sequence data are available) they may conceivably fail to detect considerable portions of the CPR taxonomic variability outside of “*Ca*. Saccharibacteria”, thus leading to a possible underestimation of some phyla and the loss of information in metagenomics studies. Indeed, a recent systematic survey analyzed the sequences from over 6,000 assembled metagenomes and evaluated 16S rRNA primers commonly used in amplicon studies. The authors observed that >70% of the bacterial clades systematically under-represented or missed in amplicon-based studies belong to CPR (Eloe-Fadrosh et al. 2016).

**Figure 1.**
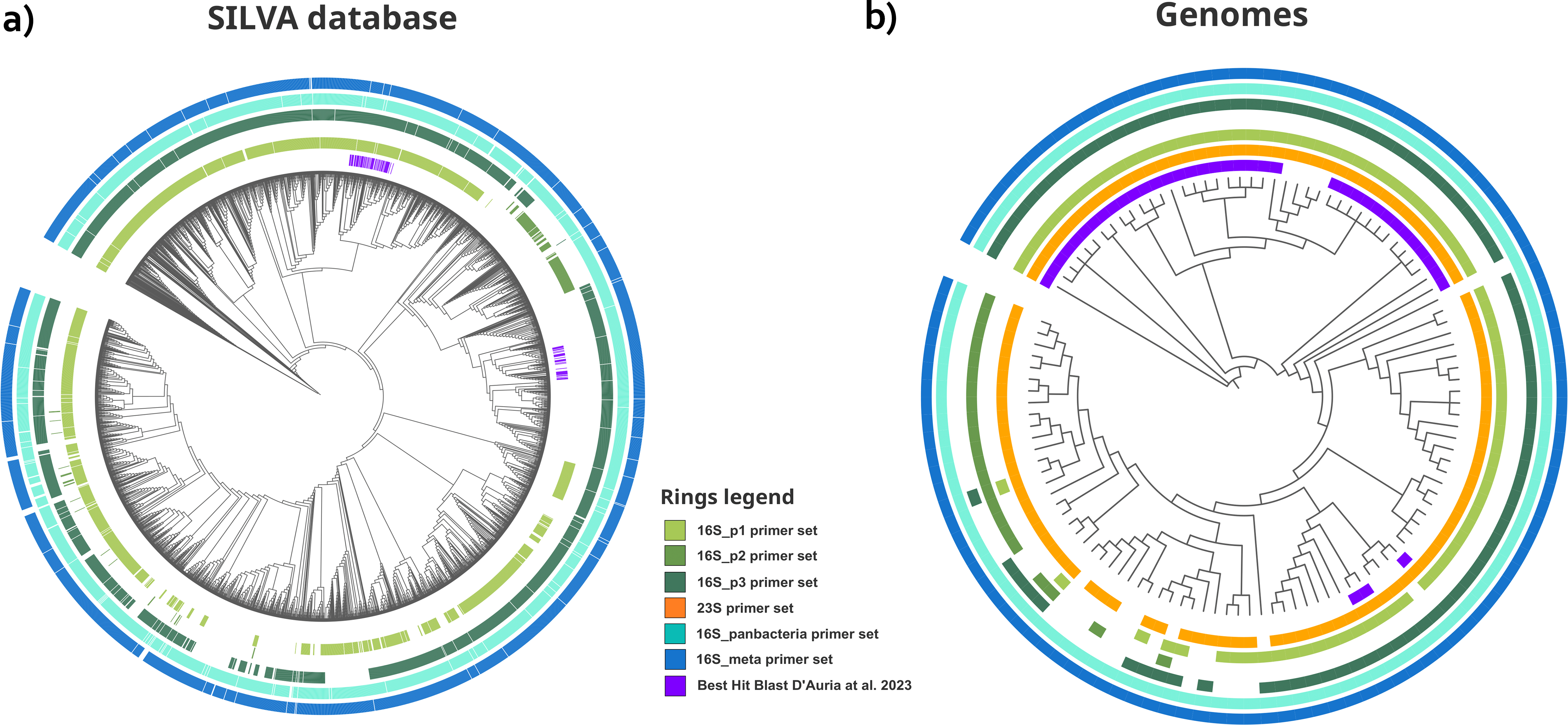
*In silico* PCR amplifications mapped on “*Candidatus* Saccharibacteria” 16S phylogenetic trees The results of *in silico* PCR amplifications are mapped on the 16S phylogenetic trees to visualize the existence of “Candidatus Saccharibacteria” lienages not amplified by qPCR primers. a) Maximum Likelihood (ML) phylogenetic tree obtained using the 16S rRNA gene sequences extracted from 114 “Candidatus Saccharibacteria” genomes harboring complete 165S and 23S genes. The inner circle (violet) shows the 16S rRNA most similar to those sequenced by D’auria et al. 2023; the second (in orange) the in silico amplification of the 23S rRNA gene on the relative genome assembly; the other circles report the *in silico* 16S rRNA gene amplifications of the primers in legend. b) ML phylogenetic tree performed on the 2,978 16S rRNA sequences annotated as belonging to “Saccharimonadia” group in the SILVA database. The circles report the the 16S rRNA most similar to those sequenced by D’auria et al. 2023 (in violet) and the the *in silico* 16S rRNA gene amplifications of the primers in legend.

Figure 1b shows the same analyses performed on the database of the 114 “*Ca*. Saccharibacteria” genomes. From the Figure it emerges that, once again, the pan- bacterial primer sets (16S_meta and 16S_panbacteria) are the most comprehensive, with a coverage of 100% (114 sequences). Among qPCR protocols, 23S was found to cover a greater portion of variability than those based on 16S. It successfully amplified 95.6% (109) of the sequences within the “*Ca.* Saccharibacteria” genome database, followed by 75.4% (86) amplified by 16S_p3 primer set, 72.8% (83) by 16S_p1, and 19.2% (22) by 16S_p2. The *in silico* PCR on 23S showed that the qPCR protocol based on this gene was able to intercept a larger portion of the “*Ca.* Saccharibacteria” taxonomic variability compared to those designed on 16S rRNA (Figure 1b). The low coverage of the 16S protocols could be attributed to the peculiar sequence and structure of the 16S rRNA gene in members of Candidate Phyla Radiation. Indeed, as stated above, it presents introns, insertions and deletions that could be an obstacle for amplification (Tsurumaki et al. 2022).

Figure 1 also maps the position, on the two phylogenetic trees, of the best hits observed for the “*Ca*. Saccharibacteria” V3-V4 16S sequences obtained by D’Auria et al. (2023). It is interesting to note that none of the sequences obtained in this paper presented a perfect match with those deposited in the two datasets. In other words, both the SILVA and genomic datasets lacked sequences whose V3-V4 portions of 16S gene were identical to those sequenced by D’auria and colleagues in their dataset, showing that the “*Candidatus* Saccharibacteria” lineages expanded in allergic children could belong to an unexplored portion within the phylum.

### 3.2. qPCR assays

The next step was to experimentally evaluate the efficiency of the selected qPCR protocols (3 based on the 16S and one on the 23S gene, Table S1) on the collection of salivary DNA previously characterized by 16S metagenomics by (D’Auria et al. 2023). In that paper, the authors found that the saliva of children suffering from food allergy, compared to matched controls, was enriched in “*Ca.* Saccharibacteria” and in sequences unresolved by the 16S metagenomics that, when phylogenetically investigated, clustered within another CPR phylum, namely “*Candidatus* Gracilibacteria”.

For each protocol and for each sample, the representation of “*Ca.* Saccharibacteria” within the bacterial community was estimated as the ratio between the “*Ca.* Saccharibacteria” quantification obtained with the specific primers (16S_p1, 16S_p2, 16S_p3 and 23S) and the total bacterial estimate obtained with the universal primer set 16S_panbacteria (Table S5). These data were then compared to the relative abundances previously obtained by the 16S metagenomics. The results of the comparisons are shown in Figure 2. The figure shows that the quantifications obtained from three out of the four protocols (23S, 16S_p1 and 16S_p2) were significantly correlated to those obtained by 16S metagenomics (linear regression, pvalue < 0.05) (Figure 2a-c). Among these protocols, only the one based on the 23S rRNA gene produced estimates comparable to the 16S metagenomics, both in terms of correlation and absolute quantification. Indeed, this protocol produced abundances not statistically different from 16S metagenomics (Mann Whitney U test, pvalue > 0.05) (Figure 2e).

**Figure 2.**
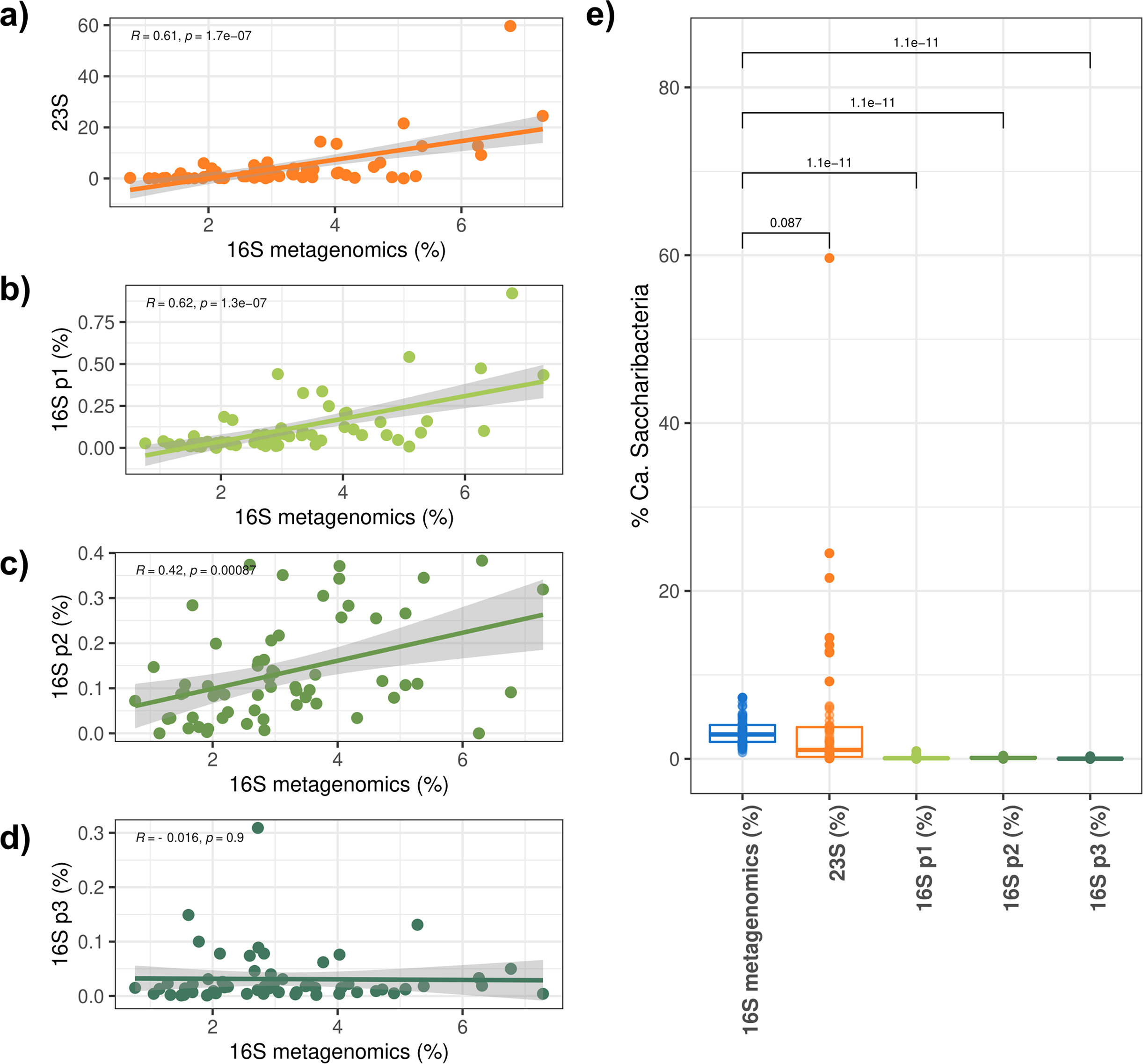
Comparison between the “*Candidatus* Saccharibacteria” quantification performed by 16S metagenomics and the four tested qPCR protocols The “*Candidatus* Saccharibacteria” relative quantification obtained by 16S metagenomics and the four qPCR protocols tested on the 61 saliva samples analysed in this study are compared. a-d) Linear regression graphs of the “*Ca.* Saccharibacteria” percentages obtained by 16S metagenomics against the 23S qPCR protocol (SacchariF-SacchariR) (a), 16S p1 (TM7314F/TM7-910R) (b), 16S p2 (Sac1031-F/Sac1218R) (c) and 16S p3 (TM7_16S_590F/TM7_16S_965R) (d). For each plot, R and p-values are reported on the top and the confidence interval is shown in gray. e) Boxplot graph of the “*Ca.* Saccharibacteria” percentages measured by 16S metagenomics and the four qPCR protocols. The median values are compared between 16S metagenomics and the other four qPCR protocols by Wilcoxon test (p-values are reported on the plot).

Instead, all the three qPCR protocols targeting the 16S gene underestimated the presence of “*Ca.* Saccharibacteria”, both in the allergic and control groups. In fact, even though two of the 16S rRNA protocols were significantly correlated with the results of the 16S metagenomics (16S_p1 and 16S_p2, see Figure 2), the absolute quantifications provided for “*Candidatus* Saccharibacteria” differed from the 16S metagenomics (and from the 23S protocol) by orders of magnitude.

Overall these results reflect the data of the *in silico* PCR conducted on the Saccharibacteria genome collection and confirm that, *in vitro* as *in silico*, the 23S protocol appears to be the most performing in terms of the portion of taxonomic diversity detected.

Another point is that the relative abundance of “*Ca.* Saccharibacteria” provided by the 16S metagenomics ranges between 0.759% and 7.286%, against a range of 0.039%- 59.665% produced by the 23S protocol (see Table S5). Thus, quantifications obtained from the 23S qPCR appear to be scattered over a much broader range than those, more flattened, provided by the 16S metagenomics. Overall, the differences between the 23S relative abundances and the 16S metagenomics ones range between -5.047% and +52.892%. Interestingly differences emerge between the two groups (controls and allergic subjects) in terms of “how much” the 23S qPCR data differ from those of the 16S metagenomics. In controls, this difference ranges within a limited interval (from - 2.455% to +10.652%) while in the allergic group it encompasses the whole interval (from -5.047% to +52.892%) (Figure S1 and Table S5).

The difference between the two quantifications was > 5% in a total of seven subjects, five allergic patients and two controls (Figure S1), thus highlighting the presence of a subset of samples, even if limited, for which the 23S qPCR protocol yielded a strongly higher quantification. For this reason, in order to exclude cross-reactions of the primers, and thus the amplification by qPCR of non-specific templates, we sequenced the 23S amplicons (see paragraph 3.3).

Among the other protocols, the best performing 16S rRNA-based qPCR was the 16S_p1. The quantifications provided by this protocol correlated with those of the 16S metagenomics but the absolute values were considerably lower. Therefore, they were not comparable in terms of absolute quantifications, clearly showing a strong underestimation of “*Candidatus* Saccharibacteria”.

There is one last important difference between the results obtained using qPCRs or 16S metagenomics. This difference is related to the increase of lineages attributable to “*Ca.* Saccharibacteria” in allergic children. While the 16S metagenomics returned a higher load of this phylum in allergic children compared to controls, these results were not confirmed by any of the tested qPCR protocols (Figure 3). This point shows very effectively how the choice to use a given technique over another can profoundly influence the final results and their interpretation in studies investigating these emerging CPR phyla and their role in the maintaining of the health status of the host. This limitation turns out to be particularly important in the case of groups such as “*Ca.* Saccharibacteria” whose role in immune-mediated diseases is increasingly evident.

**Figure 3.**
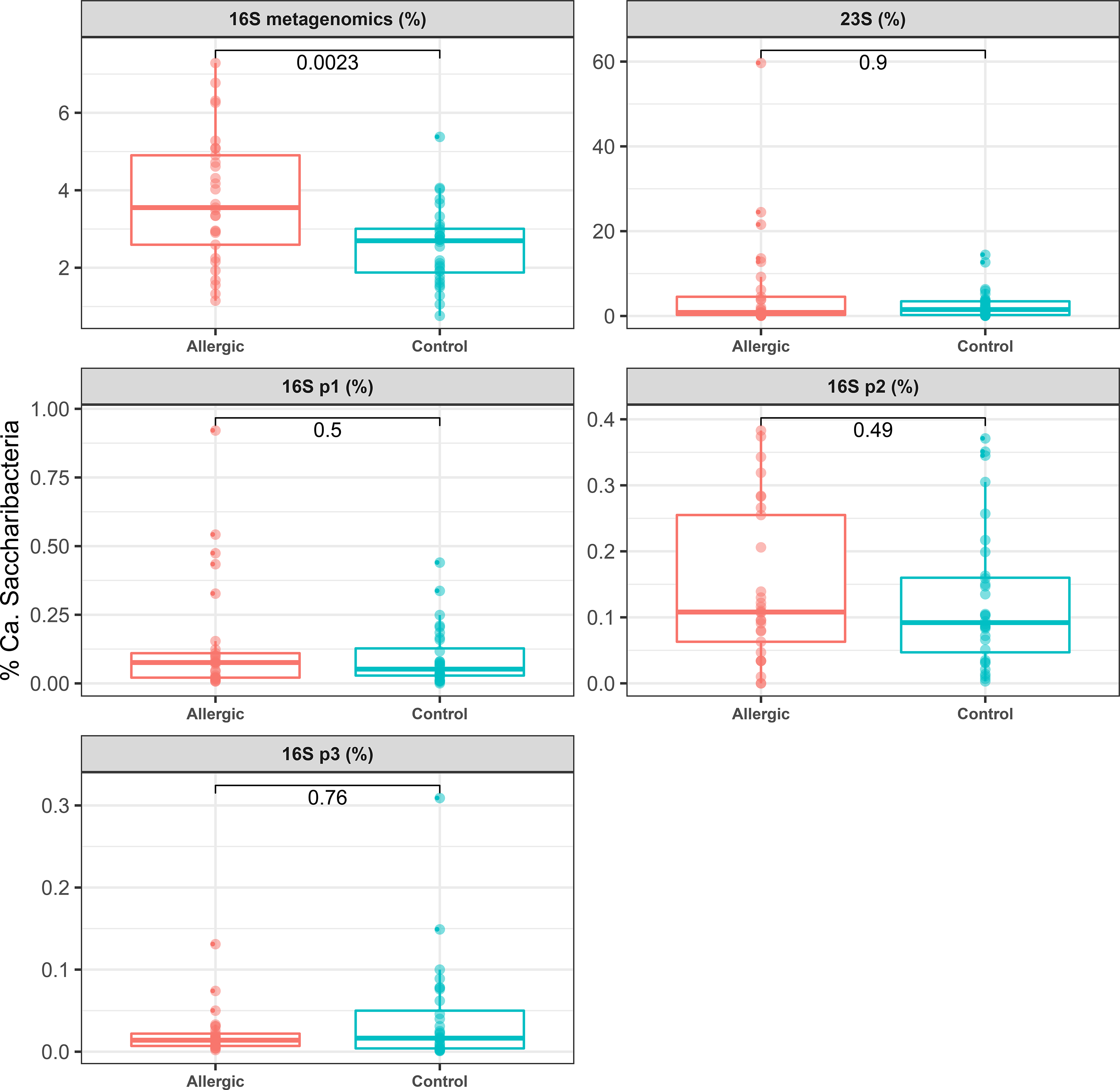
Comparison of “*Candidatus* Saccharibacteria” quantification in allergic vs control patients obtained by 16S metagenomics and four qPCR protocols a-e) Boxplots reporting the percentage of “*Candidatus* Saccharibacteria” determined by a) V3-V4 16S amplicon metagenomics, b) 23S (SacchariF-SacchariR), c) 16S p1 (TM7314F/TM7-910R), d) 16S p2 (Sac1031-F/Sac1218R) and e) 16S p3 (TM7_16S_590F/TM7_16S_965R). The values obtained from allergic vs control patients were compared using Wicoxon test and the p-values are reported on the bars.

### 3.3. 23S rRNA qPCR amplicon sequence analysis

To exclude cross-reactions and contaminations in the 23S qPCR (see above), and have direct evidence on which “*Ca*. Saccharibacteria” lineages were amplified by this protocol (the first one to target a gene other than the 16S on “*Ca*. Saccharibacteria”) amplicons from a selected subset of samples were sequenced on an Illumina platform. A total of 940,756 sequences were produced and 819,506 (87,11%) of them passed the filtering steps. The analysis grouped these sequences into a total of 11 OTUs, of which the OTU1 contains 818,910 reads, corresponding to the 99.93% of the filtered reads (Table S6). Figure S2 shows the Maximum Likelihood (ML) phylogenetic tree including the representative sequences of the 11 OTUs and the most similar sequences retrieved from the NCBI nt database. The tree topology shows that 9 out of 11 OTUs sequences (for a total of 819,498 / 819,506) clusterize within “*Candidatus* Saccharibacteria”. The remaining two OTU sequences (representing a total of 8 reads) are close to non-CPR bacteria.

These results excluded primers cross-reactions and the presence of non-specific amplicons. Therefore, the discrepancies observed with 16S metagenomics, i.e. the production, by the 23S protocol, of a broader range of quantifications, some of which are strongly higher in a subset of samples (see section “3.2. qPCR assays”), could be explained by hypothesizing the existence of “*Ca.* Saccharibacteria” lineages amplified by 23S and not by 16_meta. This point once again underlines the current lack of experimental approaches capable of detecting in a comprehensive and reproducible way the taxonomic diversity underlying “*Candidatus* Saccharibacteria” and, probably even more so, all those CPR phyla for which sequence data are even scarcer.

## 5. Conclusion

Growing evidence currently highlights the importance of having a reliable method for the detection and quantification of Candidate Phyla Radiation (CPR) members in metagenomic studies. Several papers have shown that 16S metagenomics strongly underestimates CPR and is unable to efficiently resolve their taxonomy, probably due to sequence peculiarities of this gene in CPR members. (Brown et al. 2015). It has also been estimated that >70% of bacterial clades under-represented or missed in amplicon-based microbiota surveys belong to CPR (D’Auria et al. 2023; Eloe-Fadrosh, Paez-Espino, et al. 2016). This metagenomic underestimation has several effects, particularly relevant when investigating immune-mediated diseases, considering that CPR lineages as “*Ca.* Saccharibacteria” have been experimentally observed to exert immunomodulatory roles in the human host and are enriched in several inflammatory conditions.

In recent years, several qPCR protocols targeting 16S or 23S rRNA genes have been designed for the quantification of “*Ca.* Saccharibacteria” in various environments. Four of these qPCR protocols were evaluated in this study, both *in silico* and experimentally on samples already characterized by 16S metagenomics. From the data presented in this work, we conclude that none of these experimental approaches is able to comprehensively and reproducibly detect the taxonomic diversity within “*Ca.* Saccharibacteria” and that each protocol likely introduces distortions in detection, quantification and reconstruction of taxonomic pictures. If this is the situation for the CPR phylum for which the greatest amount of sequence data has been produced, the limitations of the current protocols will likely be much greater for other CPR phyla for which sequence data are even scarcer, if not at their beginning.

On the other hand, it is becoming increasingly clear that this intriguing and ubiquitous part of the microbial world has emerging roles in important clinical and environmental processes, and that these roles have been probably greatly underestimated until now. To overcome these limitations, new experimental strategies are therefore necessary, such as the availability of new amplification targets and workflows based on amplicon sequencing. These strategies should lead to more realistic pictures of CPR abundance within bacterial communities, and of associated fluctuations (either inter-individual or associated with pathogenic processes), and allow for more efficient and precise taxonomic resolution. These premises are necessary for more targeted and systematic functional studies, to clarify their role in maintaining the health status of the host and ecological roles in the environment.

## Author Contributions

Conceptualization, F.C. and S.Pan.; formal analysis, F.C., R.N., L.S. investigation, S.Pap. and C.C.; supervision, E.D., E.P., G.Z.; writing—original draft, S.Pap. writing— review & editing, S.Pan. and F.C. All authors have read and agreed to the published version of the manuscript.

---

Figure S1. Difference in “*Candidatus* Saccharibacteria” percentages obtained by 23S qPCR and V3-V4 16S rRNA metagenomics The two histograms report the distribution of the differences between the “*Candidatus* Saccharibacteria” percentages obtained by 23S qPCR quantification and V3-V4 16S rRNA metagenomics. On the top, the distribution of the differences on the samples from allergic subjects, on the bottom those from controls. Dashed vertical lines indicate -5 and +5 percentages.

Figure S2. Phylogenetic tree of the qPCR 23S amplicon sequences On the left, the Maximum Likelihood (ML) phylogenetic tree of the sequences representative of the Operative Taxonomic Units (OTUs) of the amplicons obtained using the 23S primers (SacchariF-SacchariR) and background sequences retrieved from nt NCBI database after Blastn search. In red, the “*Candidatus* Saccharibacteria” clade and in gray the clade including sequences of non-Candidatus Phyla Radiation (CPR). The name of the OTUs and the number of sequences included in each OTU are reported on the leaves on the tree. The labels of the leaves of the sequence retrieved from the NCBI nt database are omitted.

| Protocols | In Silico PCR<br>Coverage on<br>16S rRNA<br>sequences of the<br>SILVA Database | In Silico PCR<br>Coverage on<br>16S/23S rRNA<br>sequences of Ca.<br>Saccharibacteria<br>genome dataset | In vitro PCR<br>Level of correlation<br>with V3-V4 16S<br>rRNA metagenomic |
| --- | --- | --- | --- |
| 16S_p1 | 64% | 73% |  |
| 16S_p2 | 6% | 19% | low correlated |
| 16S_p3 | 83% | 75% |  |
| 23S | / | 95% | highly correlated |
| 16S_meta | 97% | 100% | / |
| 16S_panbacteria | 97% | 100% | / |

