## Supplementary figures and images for "Unraveling and quantifying “*Candidatus* Saccharibacteria”: *in silico* and experimental evaluation of V3-V4 16S rRNA metagenomics and qPCR protocols"

### Figure S1

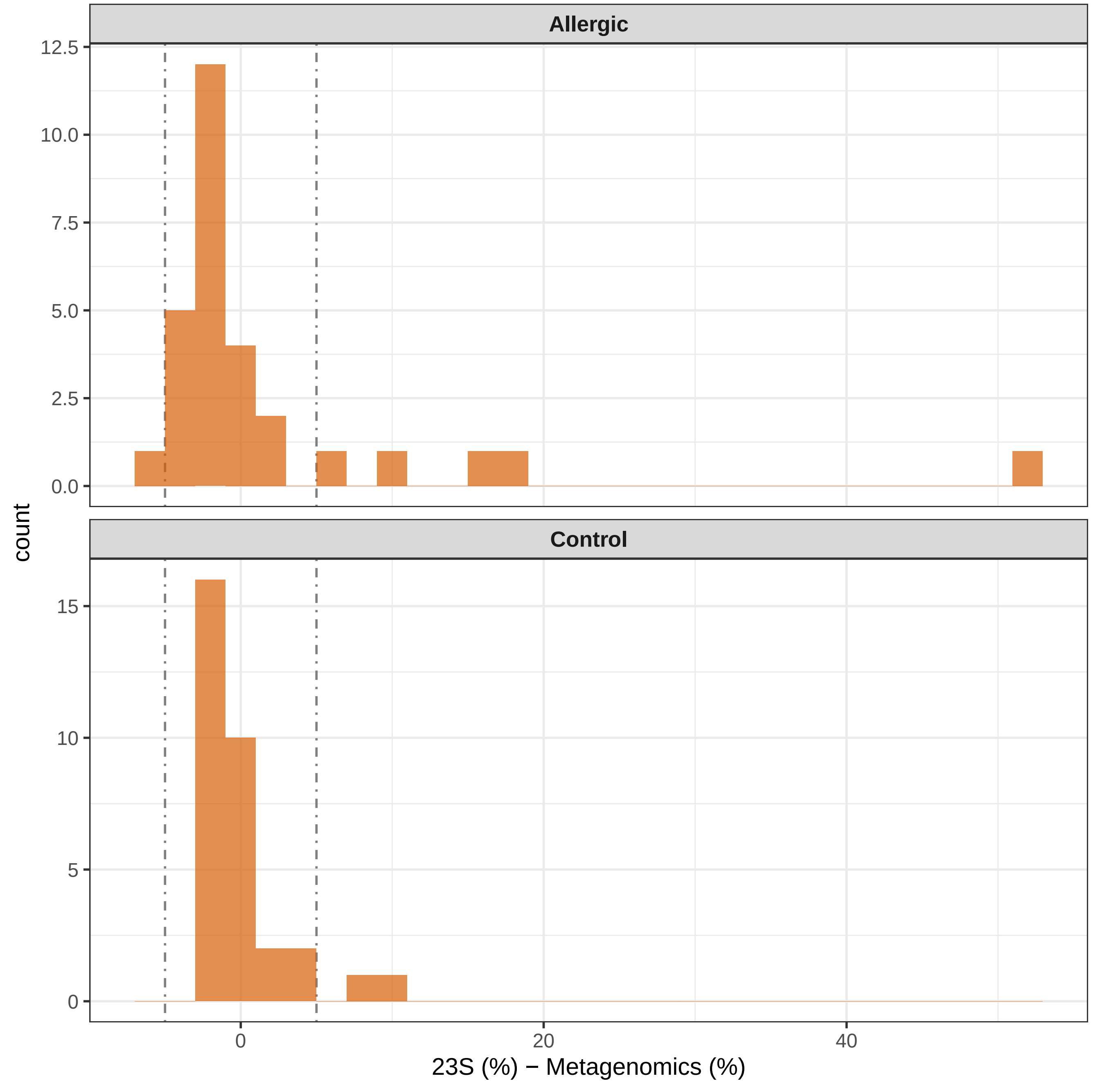

### Figure S2

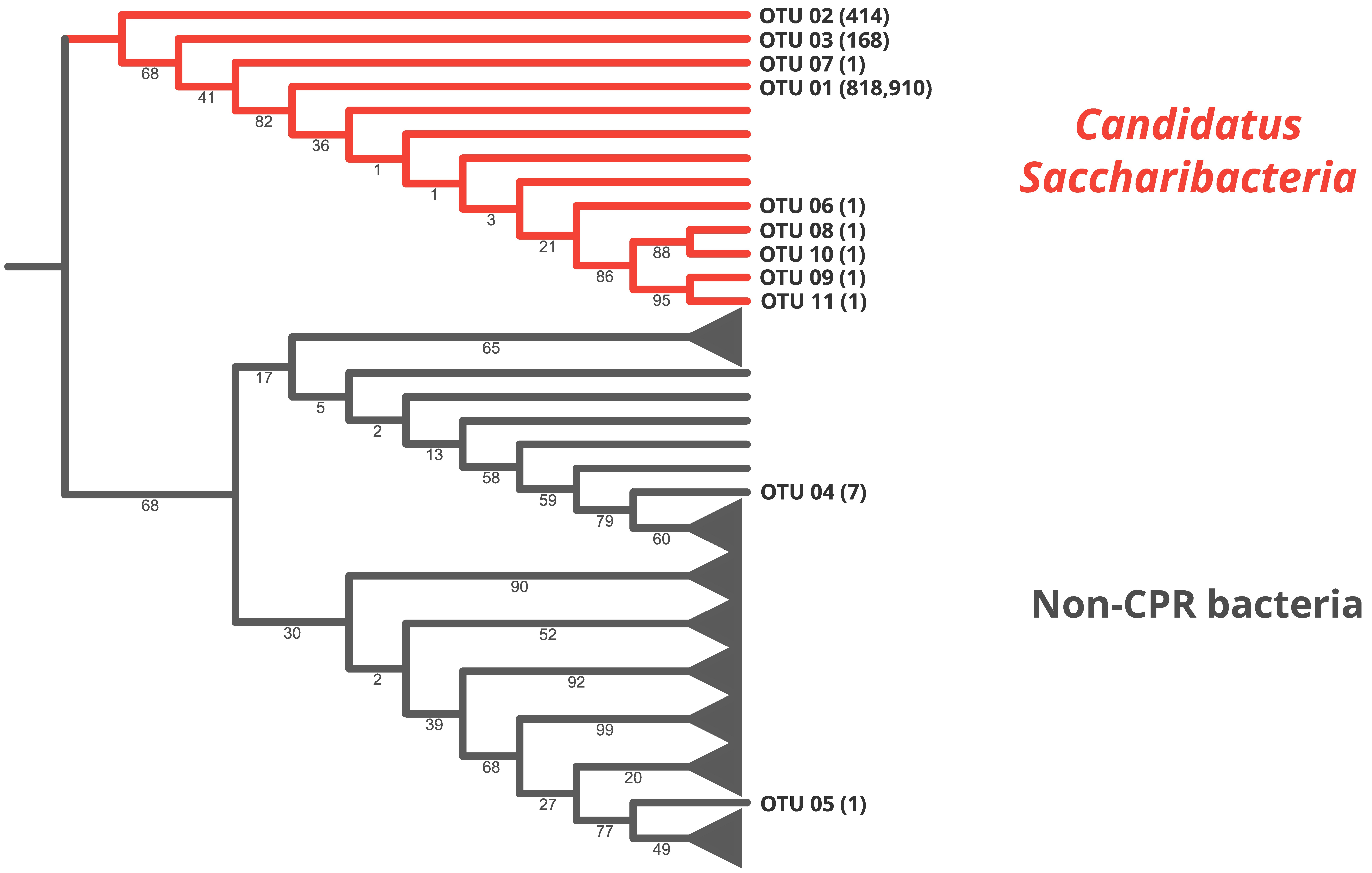
