## Supplementary material for "Unraveling and quantifying “*Candidatus* Saccharibacteria”: *in silico* and experimental evaluation of V3-V4 16S rRNA metagenomics and qPCR protocols": Table S1-S2-S3

qPCR primers tested in this work

| **Target group** | **Primer name** | **Sequence (5′–3′)** | **Protocol name** | **Amplicon length (bp)** | **Reference** |
| --- | --- | --- | --- | --- | --- |
| *Ca. Saccharibacteria* | SacchariF | GGCTTATAGCGCCCAATAG | 23S | 126 | Ibrahim et al. 2021 |
| *Ca. Saccharibacteria* | SacchariR | CGGATATAAACCGAACTGTC | 23S | 126 | Ibrahim et al. 2021 |
| *Ca. Saccharibacteria* | TM7314F | GAGAGGATGATCAGCCAG | 16S_p1 | 596 | Hugenholtz et al. 2001 |
| *Ca. Saccharibacteria* | TM7-910R | GTCCCCGTCAATTCCTTTATG | 16S_p1 | 596 | Brinig et al. 2003 |
| *Ca. Saccharibacteria* | Sac1031F | AAGAGAACTGTGCCTTCGG | 16S_p2 | 187 | Yang et al. 2015 |
| *Ca. Saccharibacteria* | Sac1218R | GCGTAAGGGAAATACTGACC | 16S_p2 | 187 | Yang et al. 2015 |
| *Ca. Saccharibacteria* | TM7_16S_590F | GWAAAGAGTWGCGTAGGYGG | 16S_p3 | 375 | Ferrari et al. 2014 |
| *Ca. Saccharibacteria* | TM7_16S_965R | WTRCTTAACGCGTTAGCTTCGCT | 16S_p3 | 375 | Ferrari et al. 2014 |
| Universal | 926F | AAACTCAAAKGAATTGACGG | 16S_panbacteria | 136 | Bacchetti De Gregoris et al. 2011 |
| Universal | 1062R | CTCACRRCACGAGCTGAC | 16S_panbacteria | 136 | Bacchetti De Gregoris et al. 2011 |
| Metagenomics | pro314F | CCTACGGGNBGCASCAG | 16S_meta | 491 | Takahashi et al 2014 |
| Metagenomics | pro805R | GACTACNVGGGTATCTAATCC | 16S_meta | 491 | Takahashi et al 2014 |

**Table S2**

Thermal profiles used in the PCR experiments

| **Primer set PCR** | **Protocol Name** | **Thermal Profile** |
| --- | --- | --- |
| SacchariF-SacchariR | 23S | 95 °C for 3 min  [95 °C for 30 s  60 °C for 30 s  72 °C for 30 s] x33  75 °C for 5 min |
| 16S_TM7_314F_910R | 16S p1 | 95 °C for 3 min  [95 °C for 30 s  62 °C for 30 s  72 °C for 30 s] x33  75 °C for 5 min |
| 16S_TM7_1031F_1218R | 16S p2 | 95 °C for 5 min  [95 °C for 15 s  61.5°C for 15 s  72 °C for 20 s] x39  75 °C for 5 min |
| 16S_TM7_590F_965R | 16S p3 | 94 °C for 5 min  [94 °C for 30 s  61 °C for 30 s  72 °C for 30 s] x34  72 °C for 5 min |
| 16S_univ_926F_1062R | 16S_panbacteria | 95 °C for 5 min  [95 °C for 15 s  61.5°C for 15 s  72 °C for 20 s] x39  75 °C for 5 min |

**Table S3**

Thermal profiles used in the qPCR experiments

| **Primer set qPCR** | **Protocol Name** | **Thermal Profile** |
| --- | --- | --- |
| SacchariF-SacchariR | 23S | 95 °C for 2 min  [95 °C for 15 s  60 °C for 7 s  72 °C for 15 s] x39 |
| 16S_TM7_314F_910R | 16S p1 | 95 °C for 3 min  [95 °C for 15 s  64 °C for 15 s  72 °C for 15 s] x39 |
| 16S_TM7_1031F_1218R | 16S p2 | 95 °C for 5 min  [95 °C for 15 s  61.5°C for 15 s  72 °C for 20 s] x39 |
| 16S_TM7_590F_965R | 16S p3 | 95 °C for 3 min  [95 °C for 15 s  61 °C for 15 s  72 °C for 15 s] x39 |
| 16S_univ_926F_1062R | 16S_panbacteria | 95 °C for 5 min  [95 °C for 15 s  61.5°C for 15 s  72 °C for 20 s] x39 |
